# “Submandibular Injection of Indoleamine 2,3-Dioxygenase Galectin-3 Fusion Protein Inhibits and Prevents Periodontal Disease Progression Over 12 Weeks”

**DOI:** 10.1101/2021.07.28.454186

**Authors:** Fernanda Rocha, Christina Graves, Ali Altitinchi, Shaheen A. Farhadi, Sabrina L. Macias, Arun Wanchoo, Gregory A. Hudalla, Benjamin G. Keselowsky, Shannon M. Wallet

## Abstract

A major challenge in the treatment of chronic inflammatory diseases including periodontal diseases is the development of therapeutics to safely and specifically direct resolution of inflammation in a localized manner. Here we demonstrate the efficacy of a novel approach that addresses this unmet clinical need. Using an enzyme fusion protein of indoleamine 2,3-dioxygenase 1 (IDO) and galectin-3 (Gal3) with tissue-anchored and thus localized immunomodulatory properties, we prevented and halted progression of induced polymicrobial periodontal disease. Efficacy was achieved with repeated treatment with IDO-Gal3 and was measured by effects on mandibular bone loss, soft tissue inflammation and local T cell plasticity. Specifically, both prophylactic and therapeutic administration of IDO-Gal3 were effective in reducing vertical mandibular bone loss and loss of mandibular bone volume and bone thickness induced by chronic polymicrobial infection. The efficacy observed is at least in part due to localized regulation of innate immune mediators such as IL-1β, IL-6 and IL-8 as well as chemoattractants including IP-10, MCP-1 and MIP-1α, while enhancing the expression of immunoregulatory cytokines such as IL-10. In addition, IDO-Gal3 treatment suppressed the induction of pathogenic CD4+Th1, CD8+Tc1, CD4+Th1/Th17, and CD8+Tc1/Tc17, while promoting the induction of homeostatic CD4+Th17, CD8+Tc17 and CD4+Tregs. Importantly, the efficacy of prophylactically and therapeutically administered IDO-Gal-3 is not dependent on bacterial burden as there were no appreciable effects on bacterial burden following either method of administration. Lastly, therapeutic rather than prophylactic administration of IDO-Gal3 is most effective in preventing mandibular bone loss through more robust innate and adaptive immune modulation. Together these and our previously published data indicate that therapeutic administration of IDO-Gal3 addresses the clinical needs for adjunctive therapeutics to treat active and/or established periodontal diseases.

## Introduction

Periodontal diseases (PD) are a group of chronic inflammatory oral and craniofacial diseases initiated by subgingival bacterial loads that result in localized destruction of both soft and hard tissues supporting the teeth. These external and internal perturbations maintain a chronic state of immune activation resulting in a vicious cycle of inflammation (1, 2). Gingivitis, the mildest yet most common form of PD, can advance to periodontitis where disease progression deepens the pocket between the gum and the tooth, more gum tissue and bone are destroyed and, without intervention, results in tooth loss (3). Current treatment of periodontitis involves mechanical removal of supra- and sub-gingival bacterial plaque and calculus (*i*.*e*., SRP) in the presence and absence of locally administered adjunctive therapies (4). However, these therapies result in modest outcomes and rarely address the vicious cycle of inflammation (3, 4). Thus, interventions aimed at resolving inflammation and restoring local tissue homeostasis in a localized manner are of significant clinical need.

Indoleamine 2,3-dioxygenase (IDO) is an immunomodulatory enzyme that plays a key role in tryptophan metabolism (an essential amino acid) along the kynurenine pathway in numerous immune and non-immune? cells including fibroblasts, epithelial cells, endothelial cells and other mononuclear inflammatory cells (5)(6). Our groups and others have demonstrated that IDO maintains homeostasis between antigen-presenting cells, T-cell differentiation and T-cell effector function (7) (8). A role for modulating IDO-driven mechanisms in curbing the chronic inflammation in PD has been implicated by several studies. Specifically, Nisapakultorn *et al*. demonstrated that IDO is expressed in human gingival epithelial cells and with fewer IDO-expressing cells found in healthy tissues compared to PD-associated tissues (6). The same group also demonstrated that IDO expression increased following stimulation with a combination of proinflammatory cytokines *in vitro* (REF). In addition, the expression of IDO in macrophages and dendritic cells is primarily induced through interferon gamma (IFN-γ)-mediated host defense in response to numerous intracellular pathogens, including etiological agents of PD such as *Porphorymonus gingivalis* (9).

Our group has developed an enzyme fusion protein using IDO and galectin-3 (Gal3), with the goal of creating a tissue-anchored and localized immunomodulatory nanomedicine (10, 11). Gal3 is a carbohydrate-binding protein, as a means to restrict enzyme diffusion via binding to tissue glycans (10). Gal3 binds N-acetylglucosamine and other β-galactoside glycans, as well as glycosaminoglycans, abundant components in mammalian tissues (12, 13), including those of the periodontium. IDO and the resultant production of kynurenine metabolites is a general regulator of inflammation in response to sterile and pathogenic inflammatory stimuli (14, 15). Our group has recently published that IDO-Gal3 not only presents a new concept of anchoring immunomodulatory enzymes for robust control of focal inflammation, but also that it is effective and translatable in multiple disease settings (manuscript under review). Specifically, we demonstrated the prophylactic and therapeutic efficacy of IDO-Gal3 in murine model of acute polymicrobial PD (manuscript under review) by showing that submandibular injection of IDO-Gal3 1) was locally retained, 2) suppressed mucosal inflammation, and 3) spared mandibular bone loss. Together these data indicated that the IDO-Gal3 nanomedicine could address the therapeutic needs in PD through localized modulation of inflammation, which prevented the progression of mandibular bone loss even after the initiation of disease. In the current study, we extend upon these findings by evaluating the prophylactic and therapeutic efficacy of IDO-Gal3 in a chronic - and therefore more translational - model of periodontal inflammation. Further, we strengthen our findings by elucidating potential mechanisms of this efficacy.

## Methods

### Murine model of periodontal disease

C57Bl/6 mice 8-10 weeks of age were subjected to oral lavage with 25 µl of 0.12% chlorhexidine gluconate for three days. Infection consisted of oral lavage with *Porphoromonas gingivalis* strain 381 (2.5×10^9^) and *Aggregatibacter actinomycetemocomitans* strain 29522 (2.5×10^9^) suspended in 25 µl 2% low viscosity carboxy-methylcellulose for 4 consecutive days, every other week for a total of 12 weeks. On the first day of each week and prior to microbial lavage, sampling of the oral environment was performed with calcium alginate swabs to determine microbial composition. One week following final lavage, the maxillae and mandibles were harvested to evaluate bone morphometric analysis, soft tissue infiltration, and/or soluble mediator expression. In addition, the submandibular lymph nodes were harvested, and resultant cell suspensions cryopreserved for flow cytometric analysis.

### Bone Morphometric Analysis

Mandibles were fixed in 4% buffered formalin for 24h, stored in 70% alcohol and scanned at 18 μm resolution using X-Ray Microtomograpy (Micro-CT) (Nano-CT V|TOME|X M 240; General Electric). NRecon Reconstruction and DataViewer Software (Micro Photonics) were used to reconstruct spatially-reoriented 3D images using anatomical landmarks; a 5.4 mm^3^ region of interest (ROI) was applied with standardized dimensions of 1.5mm (frontal) x 4.0mm (sagittal) x 0.9 mm (transversal). Anatomical landmarks were used for the standardized positioning of the ROI: frontal plane = the roof of the furcation area between mesial and distal roots of the upper first molar; sagittal plane = anterior limit was the distal aspect of the mesial root of the first molar. The thickness of the ROI on the transversal plane was set to 50 slices (900 μm) and counted towards the palatal /medial direction beginning from the image that included the center of the upper first molar in its transversal width. A standardized threshold was set to distinguish between non-mineralized and mineralized tissues where total volume and total thickness of the ROI were calculated. *Mean trabecular bone volume and thickness*: The analysis assessed the percentage of mineralized tissue (BV; BT) within the total volume/thickness (TV; TT) of the ROI and is presented as presented the BV/TV ratio (mean trabecular bone volume) or BT/TT ration (mean trabecular thickness). *Vertical bone loss*: The distance from the cento-enamel junction (CEJ) to the alveolar bone was calculated at 12 sites over three molars and averaged to calculate the average vertical bone loss in mm.

### Flow cytometry

Cryopreserved submandibular lymph node cells were thawed from liquid nitrogen at 37°C, washed, and suspended in PBS prior to incubation with a fixable Live/Dead Yellow viability dye (Life Technologies, Carlsbad, CA USA) for 10 min at RT. Following Fc receptor blocking, surface staining was performed in FACS buffer [PBS, 1% FBS, 4 mM EDTA, and antibiotics (penicillin, streptomycin, and amphotericin B)] interrogating expression of CD3, CD4, CD8, CCR6, CXCR3. Intracellular staining for FOXP3 and HELIOS was performed using FOXP3 Fix/Perm Buffer Set (BioLegend®, San Diego, CA USA). All antibodies were used at manufacturer-recommended concentrations. Fluorescence minus one or isotype controls were used as indicated. Data were acquired using a BD LSRFortessa™ flow cytometer (BD Biosciences) and analyzed using FlowJo™ Software Version 10 (Ashland, OR: Becton, Dickinson and Company; 2019). Data were normalized to 10,000 total cells collected in the lymphocyte gate and are presented as frequencies and total cell numbers.

### Soft tissue cytokine expression

Mandibles with both soft tissue and bone were subjected to bead beating at two 2 min intervals with 2 min of cooling in between using 1.0 mm diameter zirconia silica beads in cell extraction buffer prepared with a protease inhibitor cocktail and PMSF protease inhibitor to allow for dissociation and lysis of all soft tissue while leaving the hard tissues intact. MILLIPLEX® Multiplex Assays (EMD Millipore, Billerica, MA) were used to probe resulting lysates for IL-6, IL-1β, IL-10 and MCP-1 according to the manufacturer protocols. Data was acquired on a BioPlex® 200 multiplex assay system (Luminex Corporation, Austin, TX USA) running xPONENT® 3.1 software and analyzed using a 5-paramater logistic spline-curve fitting method using MILLIPLEX® Analyst 5.1 Software (MilliporeSigma, Burlington, MA USA). Data are presented as concentration [pg/ml] normalized to total protein.

### Bacterial Burden

gDNA was isolated from microbial sampling of the oral environment using a DNeasy Kit (QIAGEN, Hilden, Germany) according to manufacturer instructions. The gDNA was then probed for *P. gingivalis* 16S, *A. actinomycetemocomitans* 16S and total 16S using real time PCR. The percentage of *A. actinomycetemocomitans* 16S and *P. gingivalis* 16S within the total 16S compartment was calculated using the following formula: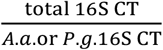

16s rRNA For: AGA GTT TGA TCC TGG CTC AG Rev: ACG GCT ACC TTG TTA CGA CTT; Pg For: CTT GAC TTC AGT GGC GGC AG; Rev: AGG GAA GAC GGT TTT CAC CA; Aa For: GTT TAG CCC TGG CCG AAG Rev: TGA CGG GCG GTG TGT ACA AGG.

### Statistical Analysis

Statistical analyses were performed using GraphPad Prism version 9.1.0 (GraphPad Software, San Diego, CA USA) where one-way ANOVA with Bonferroni’s multiple comparisons test was used to determine significance when comparing groups of three or more while Student’s *t*-test was used when comparing two groups. *P value <*0.05 was considered significant.

## Results

### Submandibular administration of IDO-Gal3 inhibits and halts mandibular bone loss following polymicrobial induced periodontal disease

In order to determine the efficacy of IDO-Gal3 in preventing alveolar bone loss, which is a hallmark of PD, IDO-Gal3 was prophylactically (IDO-Gal3-P) or therapeutically (IDO-Gal3-T) administered to C57Bl/6 mice prior to or following a 12-week regimen of polymicrobial induction of PD. As a control, uninfected mice receiving IDO-Gal3 on the same course as experimental groups (IDO-Gal3-U) were used. At 12-weeks post initial infection, mice were sacrificed, and maxillary bone volume was evaluated (**Fig. 1A**). Specifically, we measured mandibular vertical bone loss (**Fig. 1A-B**), trabecular bone volume (**Fig. 1A,C**) and trabecular thickness (**Fig. 1A,D**) and found that polymicrobial infection resulted in significant vertical bone loss as well as loss of trabecular bone volume, as expected. Prophylactic administration of IDO-Gal3 (IDO-Gal3-P) was found to reduce vertical bone loss thereby resulting in increased bone volume and thickness when compared to both control and therapeutically treated mice (IDO-Gal3-T) (**Fig. 1B-D**). Impressively, therapeutic administration of IDO-Gal3 after disease has been induced (IDO-Gal3-T), resulted in significantly better control of mandibular bone loss as demonstrated by substantially less vertical bone loss and reduction in bone volume and thickness (**Fig. 1B-D**). Together these data demonstrate that therapeutic administration of IDO-Gal3 (IDO-Gal3-T) is the most effective in preventing mandibular bone loss induced by chronic polymicrobial infection.

**Figure 1.**
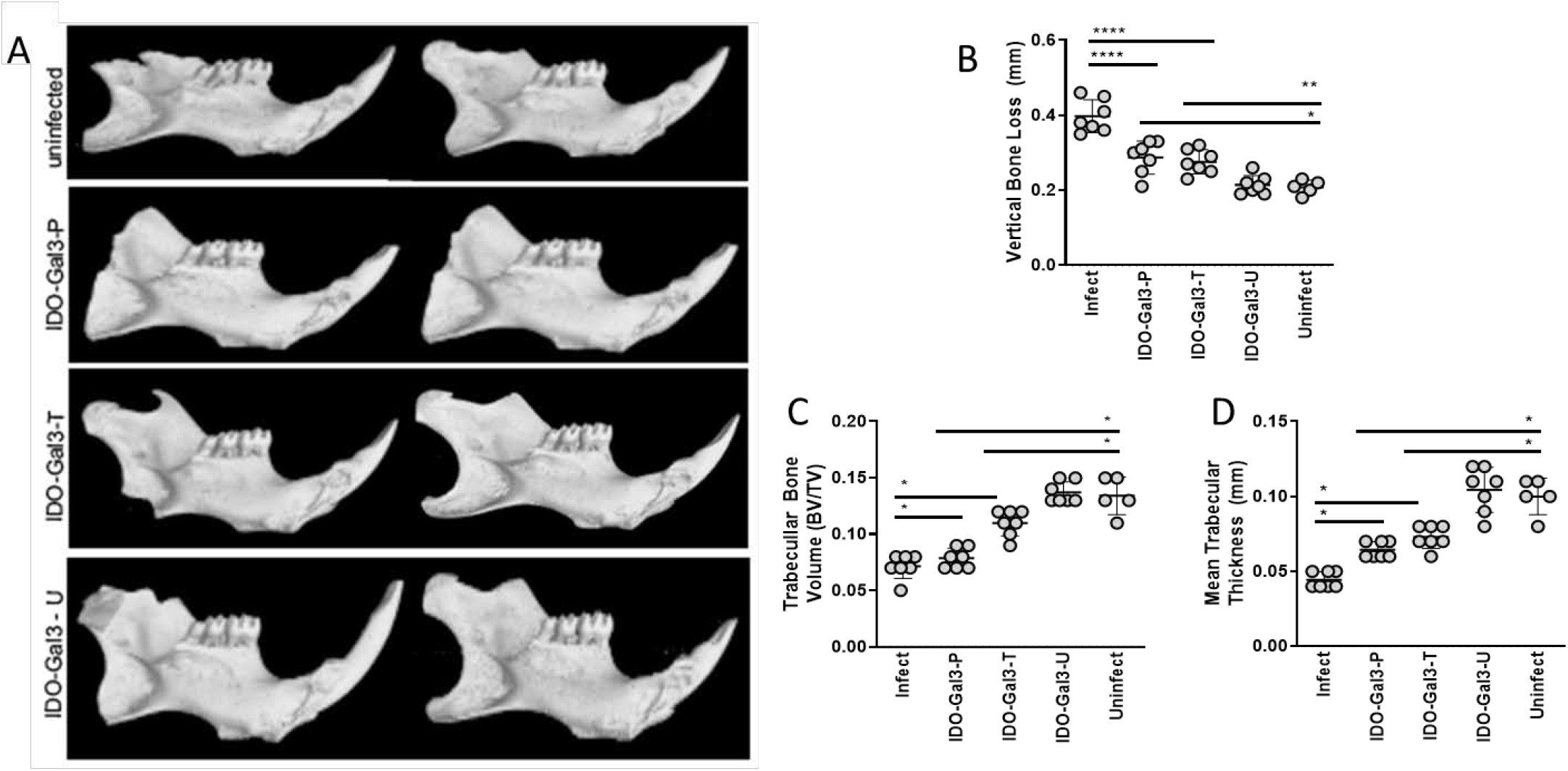
Submandibular administration of IDO-Gal3 inhibits and halts mandibular bone loss following polymicrobial induced periodontal disease. C57Bl/6 mice (n=7/group) were subjected to polymicrobial infection with 2.5×10^9^ *P. gingivalis* strain 381 and 2.5×10^9^ *A. actinomycetemocomitans* strain 29522 resuspended in 2% low viscosity carboxy-methylcellulose on 4 consecutive days (days 2 – 5), every other week for 12 weeks. IDO-Gal3 was administered prophylactically on day 1 prior to infection (IDO-Gal3-P) or therapeutically on day 6 of each week following infection (IDO-Gal3-T). Additional animals received IDO-Gal3 in the absence of polymicrobial infection (IDO-Gal3-U). **(A)** Representative images of bone morphometric analysis for each experimental group. **(B)** Vertical bone loss **(C)** Mean trabecular bone volume and **(D)** mean trabecular thickness was determined by µCT as described. Significance was determined using one-way ANOVA with Bonferroni’s correction \*\*\*\**p* <0.0001; \*\*\**p*<0.001; \*\**p*<0.01 \**p*<0.05.

### IDO-Gal3 suppresses polymicrobial induced tissue inflammation

The initiation, progression, and maintenance of PD is driven by the level of soft tissue inflammation (1, 2). Thus, the local cytokine milieu which correlated with the protection against bone loss afforded by prophylactically (IDO-Gal3-P) and therapeutically administrated IDO-Gal3 (IDO-Gal3-T) was evaluated. Compared to uninfected mice, polymicrobial infection of mice resulted in elevated soft tissue expression of the pro-inflammatory cytokines IL-1β, IL-6, and KC (**Fig. 2A-C)** as well as the chemo-attractants IP-10, MCP-1 and MIP-2 (**Fig. 2D-F**). Both prophylactic and therapeutic administration of IDO-Gal3 reduced polymicrobial induced local cytokines (**Fig. 2A-C**) and chemokines (**2D-F**). Again, therapeutic administration of IDO-Gal3 (IDO-Gal3-T) was observed to be the most effective in controlling the soft tissue inflammation to the point of restoring levels observed in the absence of polymicrobial infection. Together these data demonstrate that the efficacy of IDO-Gal3 is at least in part due to localized regulation of inflammation induced by a chronic polymicrobial infection.

**Figure 2.**
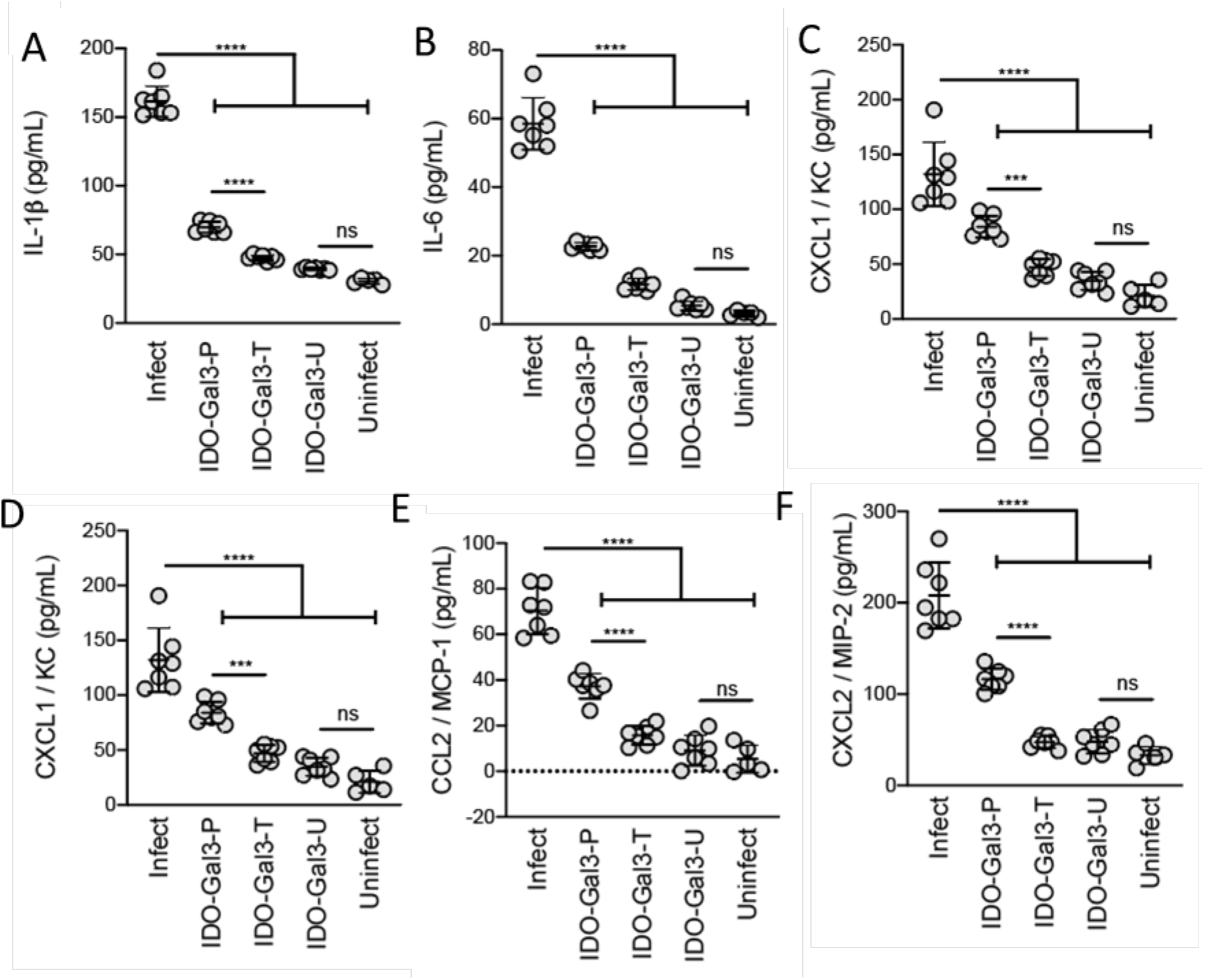
IDO-Gal3 suppresses polymicrobial induced tissue inflammation. C57Bl/6 mice (n=7/group) were subjected to polymicrobial infection with 2.5×10^9^ *P. gingivalis* strain 381 and 2.5×10^9^ *A. actinomycetemocomitans* strain 29522 resuspended in 2% low viscosity carboxy-methylcellulose on 4 consecutive days (days 2 – 5), every other week for 12 weeks. IDO-Gal3 was administered prophylactically on day 1 prior to infection (IDO-Gal3-P) or therapeutically on day 6 of each week following infection (IDO-Gal3-T). Additional animals received IDO-Gal3 in the absence of polymicrobial infection (IDO-Gal3-U). One week following the last infection, protein lysates from mandibular soft tissue was subjected to multi-parameter analysis using bioplex technology. Data are presented as pg/ml normalized to total protein. Significance was determined using one-way ANOVA with Bonferroni’s correction \*\*\*\**p* <0.0001; \*\*\**p*<0.001; \*\**p*<0.01 \**p*<0.05.

### IDO-Gal3 modulates polymicrobial induction of T cell plasticity

T cells within mucosal surfaces generally, and the periodontium specifically, are not only critical for controlling infection, but also maintaining tissue homeostasis (1, 2). Immune plasticity within T cell compartments has been described whereby the differentiation program of T cell subsets is not fixed and is highly dependent on the microenvironment in which the T cells are activated and see their cognate antigens (16). In order to determine whether IDO-Gal3-regulation of soft tissue inflammation also affected T cell plasticity, we assessed the frequency and phenotype of CD4+ T cells (Th1, Th17, Th1/17) and CD8+T cells (Tc1, Tc17, Tc1/17) within the draining lymph nodes by multicolor flow cytometry (**Fig. 3**). We found that polymicrobial infection induced the expansion or polarization of CD4+ and CD8+T cells which are CXCR3-CCR6+ (Th1 or Tc1) or CXCR3+CCR6+ (Th1/Th17 or Tc1/Tc17) at the expense of the T cells which are CXCR3+CCR6-(Th17 or Tc17) (**Fig. 3**). Infected mice which were prophylactically administered IDO-Gal3 (IDO-Gal3-P) presented with statistically fewer CXCR3-CCR6+ and CXCR3+CCR6+ cells than untreated infected mice which was concomitant with a higher frequency CXCR3+CCR6-CD4+ and CD8+ T cells. Surprisingly, infected mice which were prophylactically administered IDO-Gal3of had higher frequencies of CXCR3-CCR6+ and CXCR3+CCR6+ CD4+ and CD8+ T cells and lower frequencies of CXCR3+CCR6-CD4+ and CD8+ T cells than infected mice that received therapeutic administration of IDO-Gal3 (**Fig. 3**). Indeed, therapeutic administration of IDO-Gal3 (IDO-Gal3-T) restored frequencies of these T cell phenotypes to that observed in uninfected mice (**Fig 3**). Together these data demonstrate that the efficacy of IDO-Gal3 is at least in part due to modulation of inflammatory T cell phenotypes induced by a chronic polymicrobial infection.

**Figure 3.**
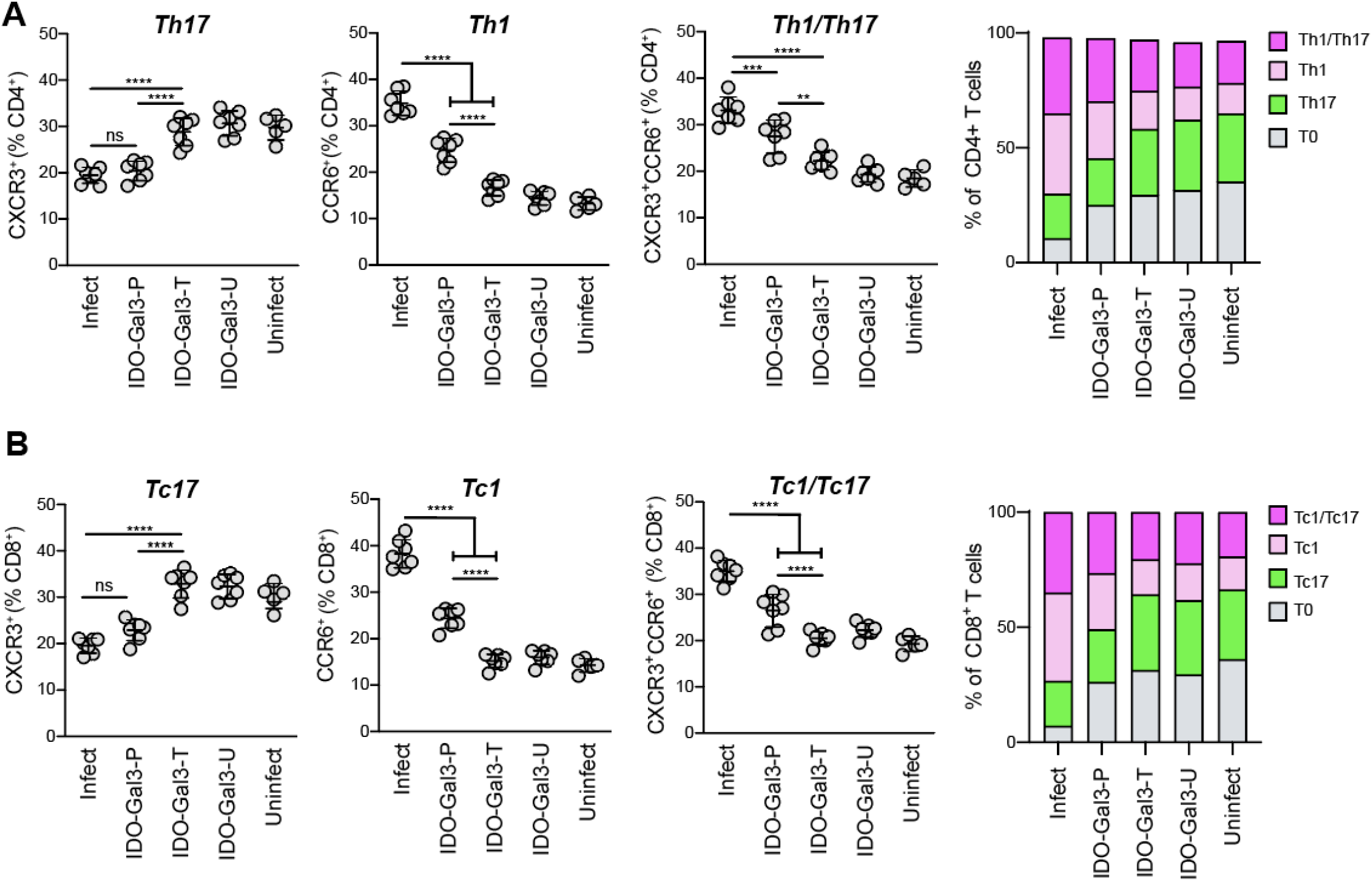
IDO-Gal3 modulates polymicrobial induction of T cell plasticity. C57Bl/6 mice (n=7/group) were subjected to polymicrobial infection with 2.5×10^9^ *P. gingivalis* strain 381 and 2.5×10^9^ *A. actinomycetemocomitans* strain 29522 resuspended in 2% low viscosity carboxy-methylcellulose on 4 consecutive days (days 2 – 5), every other week for 12 weeks. IDO-Gal3 was administered prophylactically on day 1 prior to infection (IDO-Gal3-P) or therapeutically on day 6 of each week following infection (IDO-Gal3-T). Additional animals received IDO-Gal3 in the absence of polymicrobial infection (IDO-Gal3-U). One week following the last infection, submandibular lymph nodes were subjected to multi-parameter flow cytometric analysis as previously published. Data are presented as frequencies of **(A)** CD4+CXCR3+CCR6-(Th17), CD4+CXCR3-CCR6+ (Th1), CD4+CXCR3+CCR6+ (Th1/Th17) or **(B)** CD8+CXCR3+CCR6-(Tc17), CD8+CXCR3-CCR6+ (Tc1), CD8+CXCR3+CCR6+ (Tc1/Tc17) within the CD45+ lymphocyte population. Significance was determined using one-way ANOVA with Bonferroni’s correction. \*\*\*\**p* <0.0001; *** *p* <0.001; ** *p* <0.01 * *p* <0.05.

### IDO-Gal3 promotes an immunoregulatory microenvironment needed for tissue homeostasis

Tissue innate and adaptive immune homeostasis is essential for resolution of PD, whereby the expression of soluble mediators such as IL-10 and presence of regulatory T cells (Treg) promote homeostasis (17). Thus, given the striking effect of prophylactic and therapeutically administered IDO-Gal3 on the innate and adaptive immune microenvironment, we next assessed markers of immunoregulation/homeostasis. As expected, mice exposed to a polymicrobial infection presented with significantly less mucosal IL-10 expression than uninfected mice (**Fig. 4A**) as well as a significantly lower frequency of FOXP3+ expressing CD4+ T cells within the draining lymph nodes (**Fig. 4B**). IDO-Gal3, delivered either prophylactically or therapeutically resulted in significantly higher levels of IL-10 expression in the mucosal tissues, similar to that observed in uninfected mice (**Fig 4A**). On the other hand, only mice that received IDO-Gal3 delivered therapeutically (IDO-Gal3-T) presented with an increased frequency of FOXP3+ expressing CD4+ T cells within the draining lymph nodes (**Fig. 4A**). Together these data demonstrate that IDO-Gal3 promotes tissue immune homeostasis whereby therapeutic administration is most effective.

**Figure 4.**
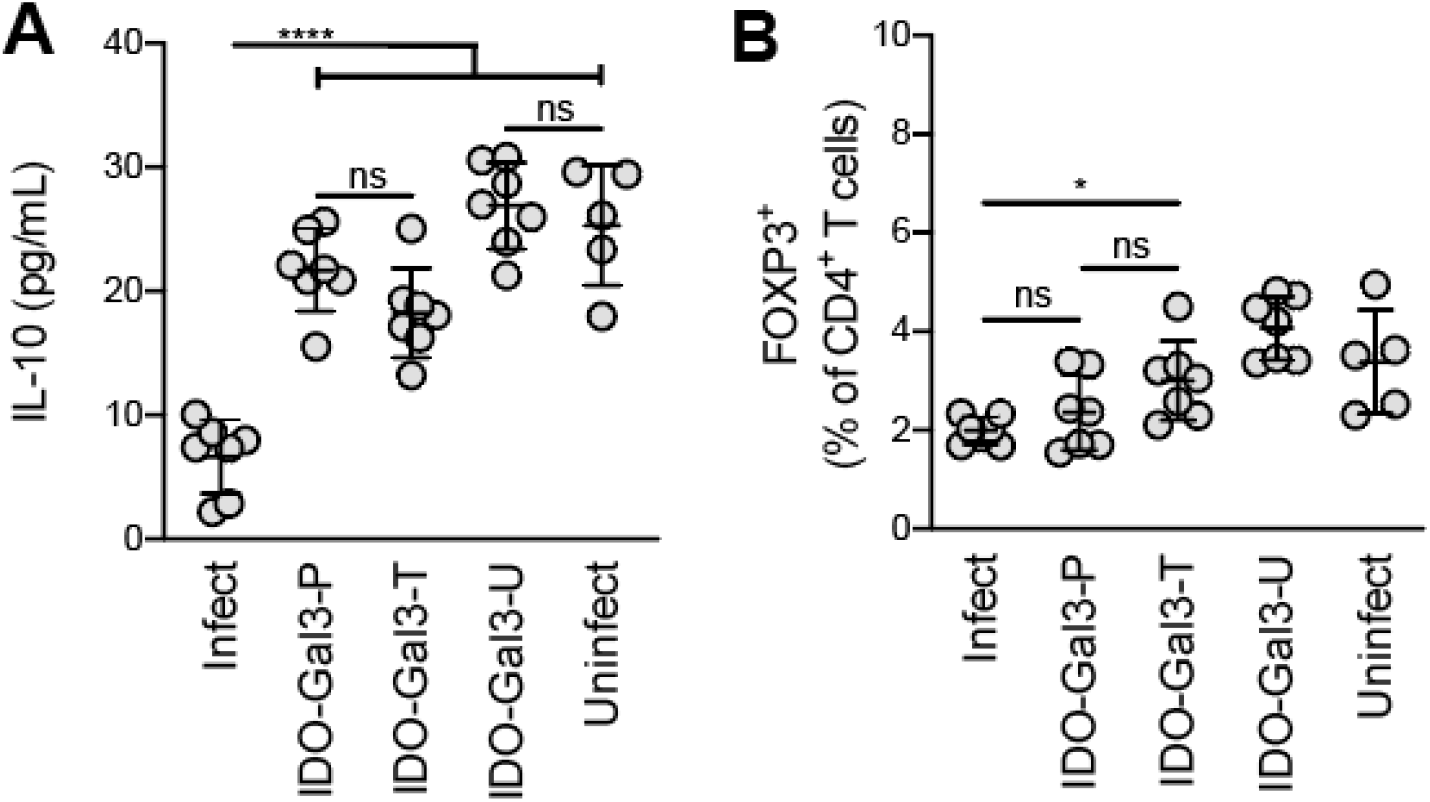
IDO-Gal3 promotes an immunoregulatory microenvironment needed for tissue homeostasis. C57Bl/6 mice (n=7/group) were subjected to polymicrobial infection with 2.5×10^9^ *P. gingivalis* strain 381 and 2.5×10^9^ *A. actinomycetemocomitans* strain 29522 resuspended in 2% low viscosity carboxy-methylcellulose on 4 consecutive days (days 2 – 5), every other week for 12 weeks. IDO-Gal3 was administered prophylactically on day 1 prior to infection (IDO-Gal3-P) or therapeutically on day 6 of each week following infection (IDO-Gal3-T). One week following the last infection, protein lysates from mandibular soft tissue and was subjected to multi-parameter analysis submandibular lymph nodes were subjected to multi-parameter flow cytometric analysis. **(A)** IL-10 expression and data are presented as pg/ml normalized to total protein. **(B)** Data presented as a frequency of CD4+FOXP3+ within the CD45+ lymphocyte population. Significance was determined using one-way ANOVA with Bonferroni’s correction. \*\*\*\**p* <0.0001; *** *p* <0.001; ** *p* <0.01 * *p* <0.05

### Efficacy of IDO-Gal3 is not dependent on suppression of bacterial infection

The PD model used in these studies is dependent on a significant bacterial burden (18) and most bacterial species, including *P. gingivalis*, utilize tryptophan in their metabolism (19). Therefore, we next assessed the impact of IDO-Gal3 on bacterial burden. Specifically, we calculated the percentage of *P. gingivalis* (**Fig 5A**) and *A. actinomycetemcomitans* (**Fig 5B**) genome relative to total bacterial genome (16S ribosome) found in the oral cavity by oral swab at the conclusion of the study. When compared to infected but untreated mice, we found that prophylactic, but not therapeutic, administration of IDO-Gal3 resulted in the enhanced recovery of *P. gingivalis* but not *A. actinomycetemcomitans* genetic material (**Fig. 5**). Despite this significance, however, this increase was less than one log-fold change, not different among other groups, and demonstrates that the efficacy of prophylactically and therapeutically administered IDO-Gal-3 is likely not dependent on bacterial burden but rather due to its immunomodulatory effects within the local immune microenvironment.

**Figure 5.**
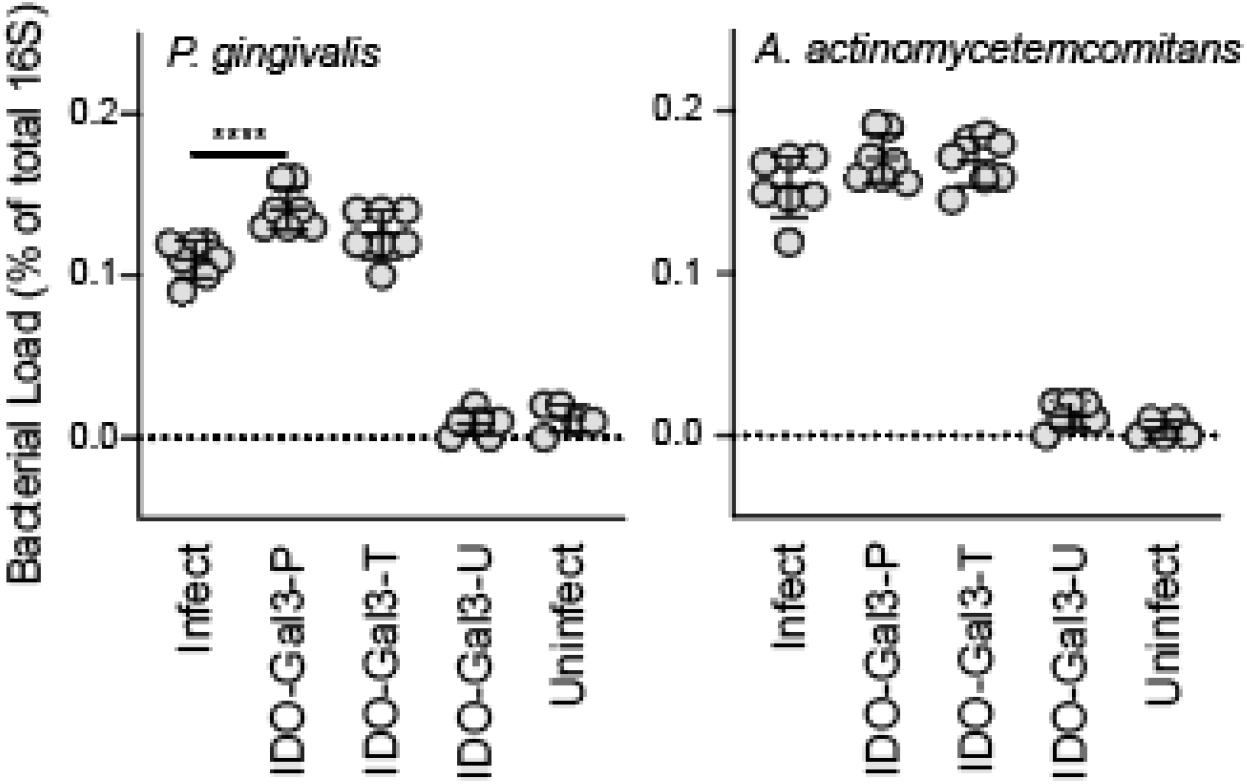
Efficacy of IDO-Gal3 is not dependent on suppression of bacterial infection. C57Bl/6 mice (n=7/group) were subjected to polymicrobial infection with 2.5×10^9^ *P. gingivalis* strain 381 and 2.5×10^9^ *A. actinomycetemocomitans* strain 29522 resuspended in 2% low viscosity carboxy-methylcellulose on 4 consecutive days (days 2 – 5), every other week for 12 weeks. IDO-Gal3 was administered prophylactically on day 1 prior to infection (IDO-Gal3-P) or therapeutically on day 6 of each week following infection (IDO-Gal3-T). One week following the last infection, RNA from oral swabs were subjected to qPCR using species specific and total 16S. Data is presented as % of total 16S. Significance was determined using one-way ANOVA with Bonferroni’s correction. \*\*\*\**p* <0.0001

## Discussion

Chronic pathologic inflammation, characterized by professional immune cell and resident tissue cell interactions, irreversibly damages tissues of the body and is an associated risk factor for a host of other diseases including cardiovascular disease, diabetes and cancer (20, 21). PD are a group of chronic inflammatory diseases whereby patients often present with these comorbidities (22). This is in part due to the external and internal perturbations which maintain a chronic state of immune activation resulting in a vicious cycle of inflammation (1, 2, 22). In the U.S., roughly 47% of the population >30 years of age suffers from PD, with the greatest incidence and severity being in adults >65 years of age (23). Thus, there is a significant clinical and public health need for novel and adjunctive treatments for PD.

A major challenge in the treatment of chronic inflammatory diseases such as PD, is the development of therapeutics to safely and specifically direct resolution of inflammation (20). Current treatment of PD involves mechanical removal of supra- and sub-gingival bacterial plaque and calculus (scaling and root planing) with or without locally administered adjunctive therapies (4). However, currently available adjunctive therapies only result in modest outcomes and rarely address the vicious cycle of inflammation.(3, 4). Anti-inflammatory small molecule drugs such as glucocorticoids are pleiotropic, non-specifically affecting numerous pathways (20). While achieving inhibition of inflammation, these drugs are accompanied by issues of toxicity, resistance, and a wide array of serious adverse effects such as infection and defective wound healing due to their broad interactions (24). Additionally, systemic administration leads to disease states such as hypertension, osteoporosis, obesity, cataracts and diabetes (24). PD is characterized by localized lesions and therefore require localized interventions. Similarly, biologic immunosuppressive drugs functioning through either cytokine blockade, cell depletion or cell surface receptor blockade provide improved specificity and can effectively modulate immune responses to halt disease progression in many, but not all patients (25). However, such treatments may also give rise to secondary infection and loss of tissue homeostasis leading to increased risk of cancer and the exacerbation of congestive heart failure and neurologic events, among other pathologies (25). Critically, each of these therapeutics require life-long use, and clinical options to fully resolve chronic inflammation and restore tissue homeostasis in a localized manner remain to be developed (26).

A major obstacle to treating the localized tissue inflammation without systemic immunosuppression is the need to accumulate therapeutics at the intended site of action. We have recently published the ability of our fusion protein IDO-Gal3 to be retained in the submandibular space for up to a week or more (manuscript under review). Gal3 is a carbohydrate-binding protein that binds N-acetyllactosamine and other β-galactoside glycans, as well as glycosaminoglycans, through its C-terminal carbohydrate recognition domain (27, 28). These glycans are highly abundant in mammalian tissues, are conserved across species, and expressed abundantly in damaged tissues (12, 13). IDO is one of the major pathways for metabolism of tryptophan in a variety of cells, including immune cells, whereby it is a critical player in establishing the balance between immunity and tolerance and ultimately in the maintenance of homeostasis (14, 15). Here we present evidence that our IDO-Gal3 fusion protein is effective in suppressing the chronic inflammation induced in our polymicrobial model of PD and restoring tissue homeostasis when given therapeutically. It is well accepted that IDO acts through *two main mechanisms*: 1) Tryptophan insufficiency via IDO activates metabolic stress sensor general control nonderepressible 2 (GCN2), inhibiting mTOR and PKC-θ for regulation of immune cell cycle and suppressive phenotypes in multiple innate and adaptive cell types (29) and 2) Kynurenine and its metabolites cause suppression and tolerance, for example through activation of the aryl hydrocarbon receptor (AhR) anti-inflammatory program, which is toxic to effector T cells and induces suppressive phenotypes in innate and adaptive cells (30-34). Additionally, kynurenine pathway production of nicotinamide adenine dinucleotide (NAD+) is regulatory in innate immunity during aging and inflammation.(35). Indeed, our data demonstrate that both the prophylactic and therapeutic administration of IDO-Gal3 results in down regulation of innate immune mediated inflammatory milieu while enhancing immunoregulation (**Fig. 2, 3**). In addition, IDO-Gal3 intervention promotes T cell phenotypes which restore and maintain local tissue homeostasis (**Fig. 3, 4**). Most importantly, the therapeuti*c* administration of IDO-Gal3 was more effective in preventing mandibular bone loss through modulation of innate and adaptive immunity. Lastly, the efficacy of IDO-Gal3 was found to be independent of bacterial burden (**Fig. 5**). These later observations are highly clinically significant considering 1) there are currently no biomarkers to predict the onset of PD and 2) not all patients achieve ideal oral hygiene. As such, adjunctive therapies which can be effective after disease has been initiated and in the absence of ideal oral hygiene are integral.

In the PD model used in these studies, efficacy was maximized with repeated IDO-Gal3 administration during the course of infection. Since the efficacy of single dosing was not investigated, we are unable to comment on the the potency of IDO-Gal3 activity within this unique bio-interface. However, our recently published data in other models of sterile and infectious inflammation indicate that a single prophylactic dose of IDO-Gal3 is sufficiently efficacious (manuscript under review). Thus, future directions include evaluating IDO-Gal3 dosage, timing, and efficacy in other models of periodontal inflammation such as peri-implantitis.

In summary, our evaluation of the prophylactic and therapeutic effect of IDO-Gal3 in a polymicrobial murine model of PD has demonstrated that submandibular IDO-Gal3 injection spares mandibular bone loss, suppresses mucosal innate inflammation, and modulates local T cell plasticity. Together these and our previously published data (manuscript under review) indicate that the IDO-Gal3 addresses the clinical needs for adjunctive therapeutics to prevent and treat PD.

## Notes

### Competing Interest Statement

The authors have declared no competing interest.

